# *Taenia solium* TAF6 and TAF9 binds to a putative Downstream Promoter Element present in *TsTBP1* gene core promoter

**DOI:** 10.1101/106997

**Authors:** Oscar Rodríguez-Lima, Ponciano García-Gutierrez, Lucía Jimenez, Angel Zarain-Herzberg, Roberto Lazarini, Abraham Landa

**Author notes:** Current address: Center for Integrative Genomics, Lausanne University, Lausanne, Switzerland (Oscar Rodríguez-Lima). Corresponding author: Abraham Landa, Ph.D. Universidad Nacional Autónoma de México. Departamento de Microbiología y Parasitología, Facultad de Medicina, Edificio A 2do piso. Ciudad Universitaria. CDMX 04510, México.

## Abstract

We have cloned and characterized the gene encoding to *Taenia solium* TATA binding protein 1 (*TsTBP1*). It spans 1481 bp and its coding region is interrupted by four introns that possess the consensus donor/acceptor sequences. It produces a protein of 238 amino acids residues, which presented all the classical motives of the TBP1. On the core promoter region we identified putative binding sites for NF1, AP-1, YY1, TAF1/TAF2 and TAF6/TAF9, and the TSS that corresponds to an A_+1_. Southern and Northern blot analysis showed that TsTBP1 is encoded by a single gene, which produce a messenger of about 1.1 kbp with a higher differential expression in adult than the larval stage. Moreover, two putative TATA boxes were identified at -97 and -69 bp (relative to the TSS), likewise a Downstream Promoter Element (DPE) was located at +27 to +31 bp. EMSA experiments did not show any component of cysticerci nuclear extracts bound to the putative TATA-box elements; in contrast TAF6 and TAF9 bind to the DPE, showing that *TsTBP1* is a TATA-less gen. On the other hand, TAF6 and TAF9 were localized on the nucleus from cells of *T. crassiceps* cysticerci bladder walls. By using the amino acid sequences of *T. solium* TAF6 (TsTAF6) and TAF9 (TsTAF9), we constructed a molecular model showing interaction between DPE-TsTAF6 and TsTAF6-TsTAF9. Finally, an *in silico* analysis of the core promoters of *Taeniidae* family genes showed that TATA-box and DPE are homologous to the mammalian elements, but not so the Inr. The low identity between TAF9 from cestodes and mammals open the possibility to use it as target to interrupt or modulate the transcription of TATA-box less genes in cestodes as a novel therapeutic strategy.

**Author summary:** Neurocysticercosis still being a health problem in developing countries. Mexico reports a prevalence of 2.5% in the total patients from National Institute of Neurology and Neurosurgery. Several efforts have been made in different fields to study *Taenia solium*, and although the Genome Project has been published and released the genomic sequence of this parasite; the transcriptional mechanisms this organism remains unexplored. We isolated and characterized *Taenia solium* TATA-Binding Protein 1 (*TsTBP1*) gene and TBP-associated factor 6 and 9 cDNAs (*TsTAF6* and *TsTAF9*, respectively). Our principal findings are: 1.- Identification of cis and trans elements in the core promoter of *TsTBP1* gene. 2.- The *TsTBP1* gene is a TATA-less promoter. 3.- Cloning of cDNAs to *TsTAF6* and *TsTAF9*, 4.- Identification of a DPE in the core promoter of *TsTBP1* gene, which interacts with TAF6 and TAF9, 5.- Construction of a molecular model that shows the interaction between DPE, TAF6, and TAF9; and 6.- A proposal of consensus sequences for TATA-box, Inr and DPE presented on *Taeniidae* family core promoters.

## Introduction

Transcription of eukaryotic protein-coding genes is regulated by RNA Polymerase II (Pol II) and a subset of protein complex known as General Transcription Factors (GTFs, TFIIA, TFIIB, TFIID, TFIIE, TFIIF and TFIIH). TFIID is the first to binds the core promoter and serves as the scaffold for assembly of the others transcription complex to position the RNA polymerase properly. TFIID is composed of the TATA-binding protein (TBP) and a group of TBP-associated factors (TAFs) which are highly conserved [1]. The core promoter is the minimum nucleotide sequence capable to direct the transcription initiation in a gene, and its cis elements are well defined in several organisms. Three cis elements are of great relevance for gene transcription: TATA-box, Initiator Element (Inr) and Downstream Promoter Element (DPE). Inr is an independent core promoter element and encompasses the Transcription Start Site (TSS, first transcribed nucleotide). The consensus for the Inr in mammalian cells is YYA_+1_NWYY, where the A_+1_ corresponds to TSS [2]. Several experiments has demonstrated that TAF1 and TAF2 binds to the Inr and form a stable complex with TBP [3–4]. TATA-box was the first core element identified and has a consensus sequence of TATAWAAR in mammalian [2]. TATA-box appears approximately at -31 to -24 bp, relative to TSS and it is recognized by TBP. It is estimated that 43% core promoters in Drosophila contain TATA-box [5] and only 32% promoters’ genes in humans [6].

The DPE was identified in *Drosophila* TATA-box-deficient promoters and it is located at +28 to +32, relatives to TSS [7], its consensus sequence is RGWYV and binds TAF6 and TAF9 [8]. It is known, by itself Inr alone in a core promoter has very low rate of transcription by Pol II, however Inr inserted downstream of six Sp1 binding sites in a synthetic core promoter, it supports high levels of transcription [9]. Recently a study demonstrated that if these three elements (TATA-box, Inr, and DPE) are presented in a synthetic promoter they enhance affinity and transcription; this was called Super Core Promoter [10]. In nature the presence and variation in the composition of these three elements in the core promoters affect their gene transcription strength [5].

TAFs are involved in basal transcription, serve as coactivators, function in promoter recognition or modify general transcription factors (GTFs) to facilitate complex assembly and transcription initiation. Biochemical and structural studies have shown that TAFs possess histone-like motifs, such as histone fold domain (HFD) that form heterodimers as TAF6-TAF9 that interact with other TAF-complex to form TFIID. It is known TAF6 and TAF9 bind to TBP and TAF1. However, there is few information about TAFs and its function [11–12]. Noteworthy, TAF9 and TAF6 have been only reported in Cestodes in genomic projects [13].

To date, there are few studies about *Taeniidae* family transcription machinery. Recently, we reported a cDNA that encodes for *Taenia solium* TBP1 (TsTBP1), the protein bind to the TATA-box of the core promoter of the Ts2-CysPrx (TsPrx) and Actin 5 (pAT5) genes and it is localized in nucleus of cysticerci [14].

In this work, we cloned and characterized the gene encoding for TATA binding protein 1 of *Taenia solium* (*TsTBP1*), and showed the existence of cis and trans elements on its core promoter, the presence of TAF6 and TAF9 in the nucleus of cysticerci, and its interaction with the DPE of *TsTBP1*. Additionally, we showed TATA-box and DPE on core promoter of *Taeniidae* family genes are homologous to the mammalian elements, but not so the Inr element.

## Materials and Methods

### Biological materials

Cysticerci from *T. solium* were dissected from naturally infected porks obtained from slaughterhouses in Mexico City, and *T. crassiceps* WFU strain cysticerci were obtained from experimentally infected mice, maintained in the Animal Facility Center from School of Medicine, National Autonomous University of México. Cysticerci were washed three times with sterile phosphate-buffered saline (PBS) and stored at −70°C, until use. Some *T. crassiceps* cysticerci were used for confocal microscopy. The *T. solium* adults were obtained from small intestine of experimental infected hamsters as previously reported [15].

### Ethics Statement

Animals were euthanized according to the Official Mexican Norms: NOM-062-ZOO-1999 for production, care and use of laboratory animals, and NOM-033-SAG/ZOO-2014, methods for killing domestic and wild animals. The research protocol was approved by the Research and Ethics Committee of the School of Medicine, National University of México (007-2012).

### Gene isolation

Preparation of *T. solium* genomic DNA and screening of 100,000 clones from a λZAPII genomic DNA library of *T. solium* cysticerci were carried out as previously reported [16], using a full length cDNA probe encoding for TsTBP1 [13]. The clones obtained (~1.5 kbp) were amplified using a primer from λZAPII vector, purified, and cloned into pCRII vector (Invitrogen, Carlsbad, CA) following manufacturer’s directions. Plasmids were sequenced on an automated DNA sequencer ABI Prism model 373 (Applied Biosystems, Grand Island, NY). PCGENE and Clustal X were used for the analysis of nucleotide sequences and multiple alignments, and Patch 1.0 BioBase server (http://www.gene-regulation.com) was used for the core promoter element analysis.

### Transcription start site determination

*Taenia solium* total RNA was prepared with TRIzol (Invitrogen) and used as template for TSS determination using the Smart 5’-RACE a cDNA Amplification Kit (Clontech, Mountain View, CA). 5’-RACE fragments were amplified by PCR using reverse primer TBPRE (inner, 5’-TCTTATCTCAAGATTTACTGTACACAC-3’), TBPRE2 (outer, 5’-CACAATGTTTTGTAGCTGTGGCTGAGG-3’) and forward primer SMARTII (5’-AAGCAGTGGTATCAACGCAGAGTACGCGGG-3’) following the manufacturer’s directions. The resulting amplified band was cloned into pCRII (Invitrogen) and sequenced as above.

### Southern and Northern blot analysis

Southern blot was carried out using 10 µg of genomic DNA digested separately with *Hind*III, *BamH*I, *EcoR*I, and *Bgl*II. Digested DNA was resolved on a 1% agarose gel and blotted on a nylon membrane (Amersham, Pittsburgh, PA). For Northern blot, total larval RNA (10 µg) from *T. solium* cysticerci was resolved by electrophoresis on a 1% agarose gel with formaldehyde and blotted to a nylon membrane. Prehybridization and hybridization were performed according to Sambrook et al [17]. As a probe we used the full cDNA coding for TsTBP1.

### Relative transcription of TsTBP1 measured by Real-time PCR

For the Real-time PCR, 3 µg of total RNA from larval and adult stage from *T. solium* was reverse-transcribed to cDNA using SMARTScribe Reverse Transcriptase (Clontech) according to manufacturer’s instructions. Thus, 200 ng of cDNA was used in a final reaction volume of 10 µl per reaction. We used the designed primers TBP-X1 (5’-ATGCAGCCAACCCCCATCAATCAG-3’) and TBP-X2 (5’-TTAGCCAGTAAGTGCTGG-3’). Likewise, *T. solium* Cu/ZnSOD amplification was done with the primers SOD-6 (5’-AAGCACGGCTTTCACGTCC-3’) and SOD-2 (5’-ACGACCCCCAGCGTTGCC-3’). The reactions were performed using the LightCycler 480 SYBR Green I Master in the LightCycler 480 System (Roche). The PCR scheme used was 95°C for 10 min, and then 40 cycles of 95°C for 15 sec and 52°C for 1 min and 72°C for 30 sec. The mRNA levels of TsTBP1 were normalized to the Cu/ZnSOD. The relative amounts of mRNA were calculated using the comparative CT method.

### cDNAs isolation to *T. solium* TAF6 (TsTAF6) and TAF 9 (TsTAF9)

Fragments of cDNA encoding for TsTAF6 and TsTAF9 were produced by PCR using *T. solium* larval cDNA as template and two oligonucleotides designed from initiation and end of each TAF6 and TAF9 sequences reported in the genome of *T. solium* project [13], The primer sequences were: TsTAF6F (5’-AGTGTTATGTTTTCGGAAGAGC-3’), TsTAF6R (5’-CTAGTGGGAGACATTGAGGG-3’), TsTAF9F (5’-ACTCTAATGAAGTTGCGTGCTG-3’) and TsTAF9R (5’-AGACGTTTAATTGCTGTGATTCC-3’). PCR conditions were: denaturation for 3 min at 95°C, followed by 30 cycles of 1 min at 95°C, 1 min at 50°C and 2 min at 72°C, and a final extension step by 5 min at 72°C. The amplified products were cloned into the pCRII vector (Invitrogen), sequenced and analyzed with PCGENE program.

### Localization of TsTAF6 and TsTAF9 by immunofluorescence

Cysticerci from *T. crassiceps* were embedded in Tissue-Freezing Medium (Triangle Biomedical Science, Durham, NC), frozen in liquid nitrogen, and stored at -70°C. Frozen of 6 to 8 µm thick sections were prepared and fixed with 4% paraformaldehyde in PBS. Samples were permeabilized with 0.01% (v/v) Triton-X 100 for 30 min, and blocked with 3% (w/v) BSA in PBS for 30 min; sections were incubated overnight with rabbit anti-TAF6 or mouse anti-TAF9 (1:100 dilution; Santa Cruz Biotechnology, Dallas, TX) antibodies. Sections were rinsed three times with PBS and incubated 60 min at room temperature with Alexa 488-conjugated anti-rabbit IgG or Alexa 568-conjugated anti-mouse IgG (diluted 1:200 in PBS-3% BSA, Life Technologies, Grand Island, NY). Sections were rinsed three times with PBS and incubated 5 min at room temperature with 4′,6-diamidino-2-phenylindole (DAPI). Normal rabbit IgG was used as control at the same concentration of the first antibody. Sections were rinsed as before and mounted on a glycerol-PBS solution (9:1). Single plane images were obtained with a confocal microscope LSM-META-Zeiss Axioplan 2. Co-localization analysis was performed using the ZEN 2010 program version 6.0 (Carl Zeiss, Pleasanton, CA).

### Electrophoretic Mobility Shift Assays (EMSA)

Nuclear extracts for EMSA was prepared as previously described [14]. To generate double-stranded probes complementary oligonucleotides were mixed at a 1:1 molar ratio and heated to 95°C for 5 min and gradually cooled to room temperature. Double-stranded probes (Table 1) were labeled with [γ-32P]ATP (Perkin Elmer, Boston, MA) using T4 polynucleotide kinase (Invitrogen). Binding reactions were performed by pre incubating at room temperature: 17.5 fmol of each labeled probe, 1µg of poly(dI-dC) and 5 or 10µg of nuclear extracts in 1X Binding buffer (20% glycerol; 2.5 mM EDTA; 5 mM MgCl_2_; 250 mM NaCl; 50 mM Tris-HCl; 2.5 mM DDT). For competition, 25 and 50-fold excess unlabeled double-stranded probe was added to the binding reaction. The reaction was incubated 30 min at room temperature. For super shift assay, 1 µg of rabbit anti-TAF6 or mouse anti-TAF9 antibodies (Santa Cruz Biotechnology, Dallas, TX) was added to the reaction 30 min after nuclear extract and incubated 30 min at room temperature. Reactions were finished with addition of gel-loading buffer. The complex was separated on a nondenaturing 5% polyacrylamide gel and visualized by autoradiography of dried gel.

**Table 1.**
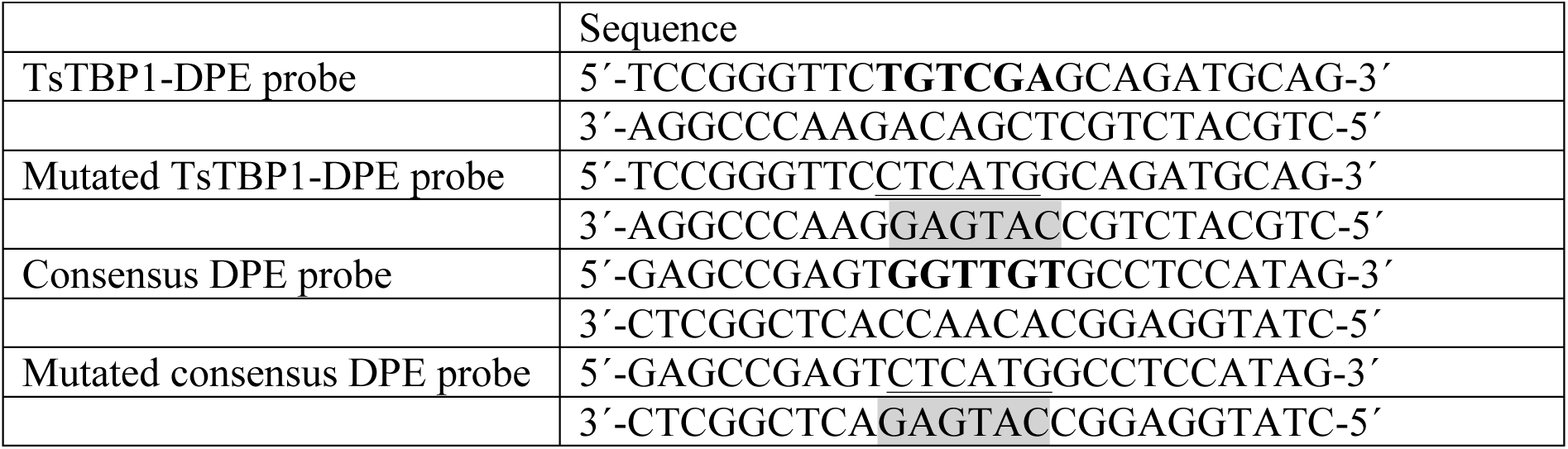
Double-stranded DNA (dsDNA) probes used for EMSA. Downstream promoter element (DPE) is in bold letters; mutated sequences are *underlined* and *bold*, and complementary mutated sequence are *highlighted* in gray.

### Molecular modeling of TsTAF6-DPE complex

TsTAF6, TsTAF9, and DPE molecular models were constructed by threading or homology modeling according to the previous protocol [14] using the sequences reported here. 3-D models were equilibrated by Molecular Dynamics (MD) simulations for 20 ns. MD simulations were performed using GROMACS 5.0.6 software [18, 19] with AMBER03 force field [18]. The side chain ionization states of TsTAF6 and TsTAF9 at pH 7.4 were established according to pKa values estimated by the PROPKA subroutine in PDB2PQR server [20]. In a first instance, models of TsTAF6 and DPE was placed into the center of a periodic dodecahedral box at distance of 30 Å between the positive patch side of TsTAF6 and the longitudinal side of DPE without any special orientation for this latter. Other five systems consist of the DPE-probe molecule placed along of the rest of Cartesian axes respect to the center of mass of TAF6C such that any two atoms from different molecules are not closer than 30 Å at the initial configuration.

The edges of the box were fixed at 20 Å from any molecules. A total of 33 Na^+^ and 36 Cl^¯^ ions were added to neutralize the net charge of the system and to fix an ionic force equal to 0.10 mol L^-1^, and 79,658 TIP3P water molecules to fill the box. The system has subjected to an energy minimization, and then a 100 ps of thermal equilibration under the position restrains to all heavy atoms by a harmonic force constant of 1000 kJ mol-1 nm-1- MD simulation was performed using an NPT ensemble at 365.15 K and 1.0 bar for 100 ns. A LINCS algorithm [21] was applied to constrain the length of all covalent bonds. The time step was 2 fs. A cutoff of 1.2 nm was applied for short-range electrostatic and van der Waals interactions, while the long-range electrostatic forces were treated using the particle mesh Ewald method [22].

### Modeling of TsTAF6-DPE-TsTAF9 system

In a first instance, pyDock [23] Web server was used to construct the tertiary molecular system TsTAF6-DPE-TsTAF9 by using the conformation of the TsTAF9 corresponding to the last frame in the dynamic simulation of TsTAF6-DPE system, and the TsTAF9 structure obtained by homology modeling. One consensus model obtained by clustering in docking process was selected as a starting point to assembly the TsTAF6-TsTAF9 system. Then we superpose the DPE coordinates according the previous TsTAF6-DPE model, allowed us to get the TsTAF6-DPE-TsTAF9 tertiary system. The same MD protocol was applied to this tertiary system, during 50 ns. The Adaptative Poisson-Boltzmann Solver (APBS) program [24] was used within PyMOL to display the results of the calculations as an electrostatic potential molecular surface. The ionic strength was set to 100 mM of NaCl.

## Results and discussion

### Genomic analysis of *TsTBP1*

*TsTBP1* sequence spans 1481 bp (Fig.1), and encodes a protein of 238 residues with a molecular mass of 27.6 kDa that contain all the characteristic motives of the TBP1, as previously reported [14]. The proximal promoter analysis reveals the presence of putative binding sites for GTFs, such as TBP at -97 bp and -69 bp (TATA-box), TAF6/TAF9 at +27 bp (DPE), TAF1/TAF2 at -2 bp (Inr), YY1 at -119 bp, AP-1 at -57 bp and NF1 at -192 bp, and two CCAAT box at -173 and -167 bp. A 5’-RACE let us identify the transcription start site (TSS, Figure 2) which corresponds to an adenine (A_+1_) located within the initiator (Inr, GCATCCT) and mapped at 36 bp upstream of the translation start codon (ATG), it matches with the first nucleotide of the cDNA clone of the encoded TsTBP1 reported previously [14]. Additionally, the structural coding region has five exons split by four introns (Intron I: 76 bp length, from +192 to +267; Intron II: 208 pb length, from +512 to +719; Intron III: 69 pb length, from +825 to +894; Intron IV: 124 pb length, from +991 to +1114 bp relatives to TSS) that present the NGT-AGN donor-acceptor sites. It showed a putative U1 recognition sequence (Intron I: <u>GT</u>AAGC; Intron II: <u>GT</u>AAGA; Intron III: <u>GT</u>TTCG; Intron IV: ACAG<u>GT</u>, showing the donor underlined site), and a pyrimidine rich tract for U2 Associated Factor (U2AF) binding site (Intron I: TGTCTCTTTTACTTTC<u>AG</u>; Intron II: TTTGCTTCCCTGCTTT<u>AG</u>; Intron III: GGAGATTTTCTTCTTT<u>AG</u>; Intron IV: <u>AG</u>ACACTTTCCATACTTG, showing the acceptor underlined site). The presence of all the elements previously mentioned suggests that *TsTBP1* possess the classic structure of a housekeeping gene, as we support later.

**Figure 1.**
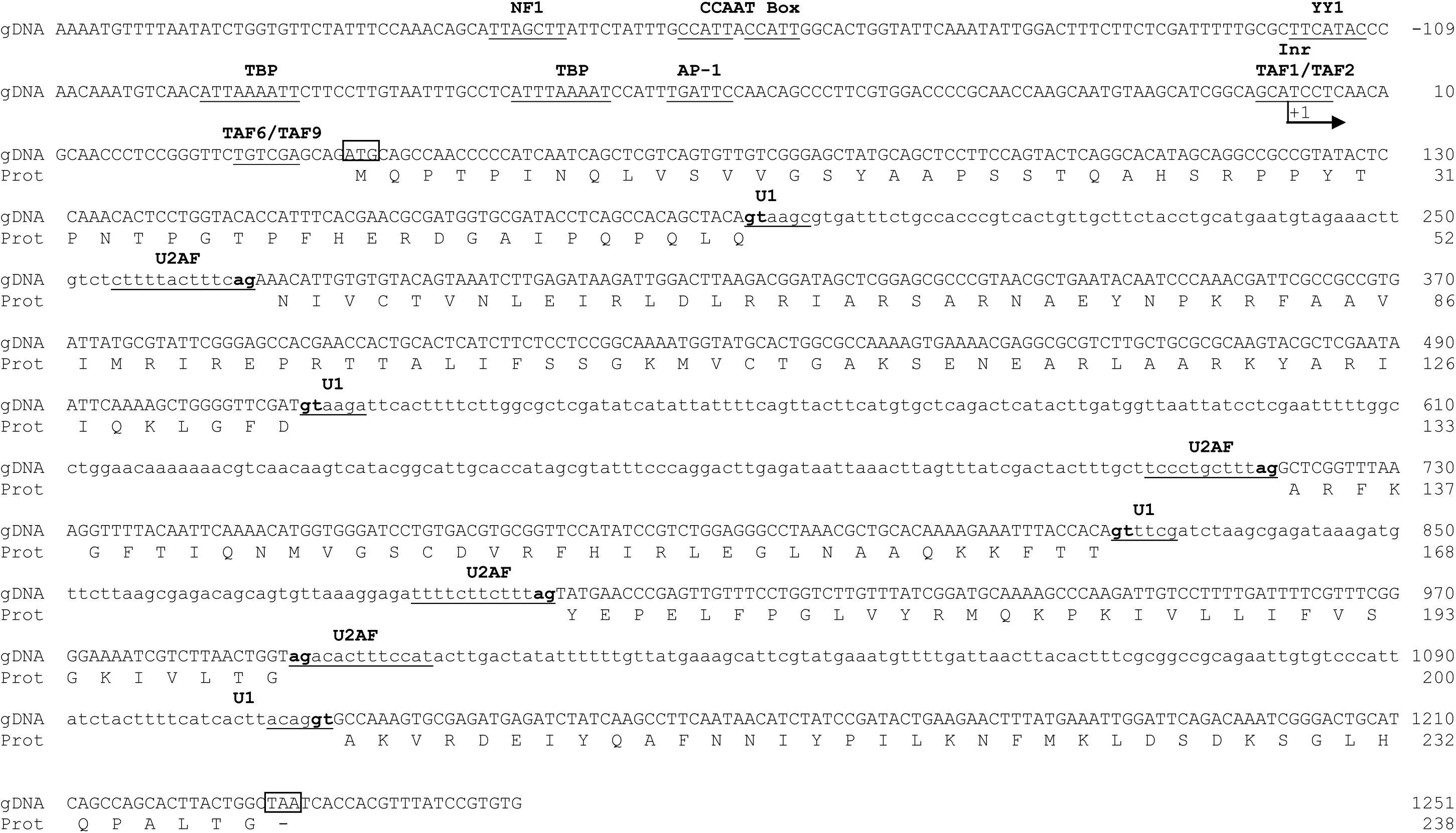
Genomic structure of TsTBP1 (GenBank accession number: KX905241) Transcription Start Site (TSS) corresponds to an A+1 marked by an arrow. Putative transcription factors binding sites are written above their underlined target sequence. Start (ATG) and Stop (TAA) codons are in a boxed. Donor and acceptor intron sequence (gt/../ag) are in bold letters. Putative U1 and U2AF splicing binding sites are underlined. Introns are written in small letters. Numbers at the right correspond to nucleotides and the amino acids translated of this gene.

**Figure 2.**
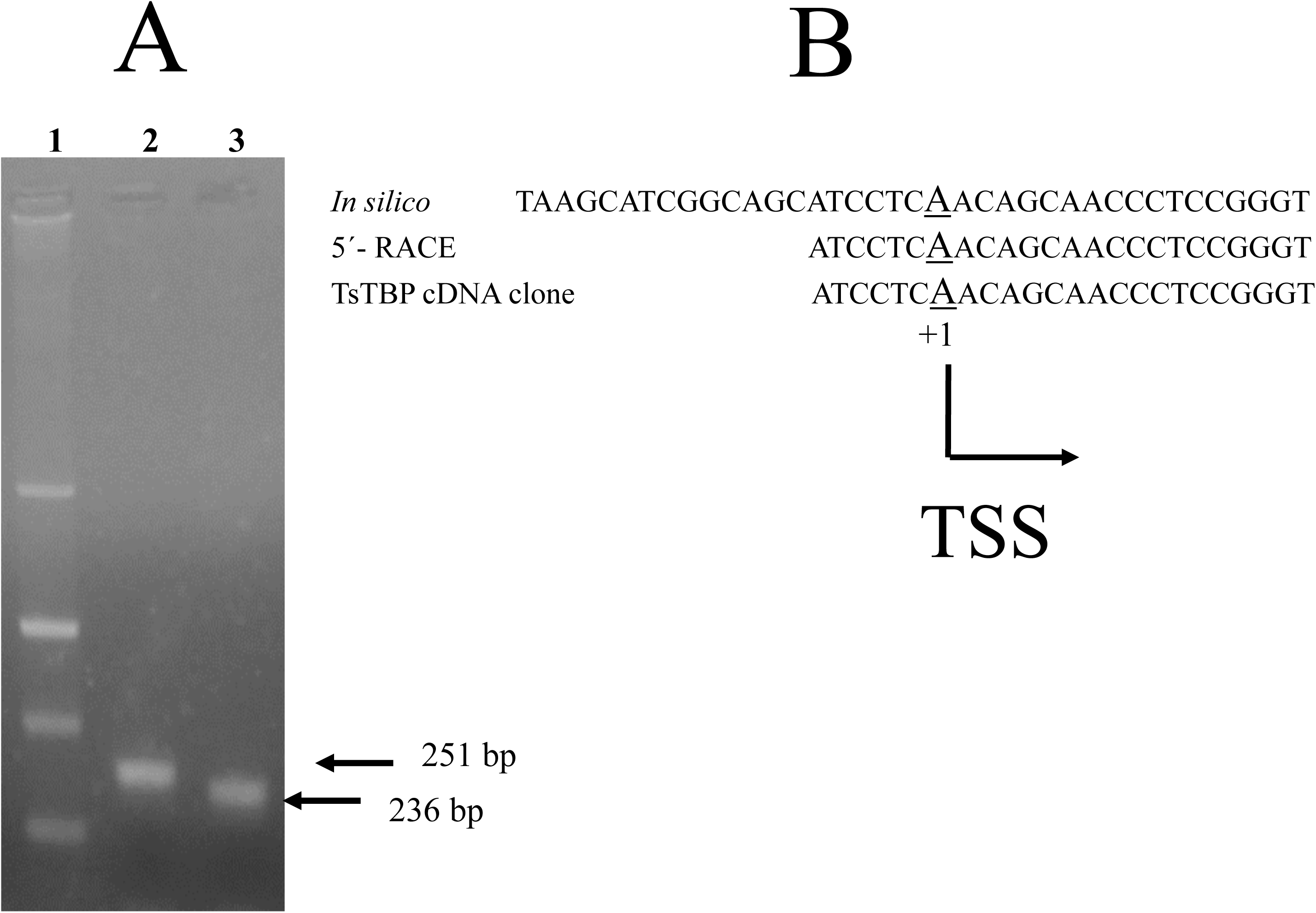
Identification of transcription start (TSS) site by 5’-RACE. A) Agarose gel shows the amplification with a T. solium cDNA ends-5’ and SMART primer versus: 2) outer primer that gives a band of 251 bp, and 3) inner primer that gives a band of 236 bp. B) Alignment of nucleotides showing the TSS from TsTBP1 obtained by: In silico analysis, experimental 5’-RACE and the cDNA full-length clone (TsTBP1 cDNA clone) reported (14).

*TsTBP1* has four small introns with different lengths, and it is known that the number of introns varies between species. It is noteworthy that *O. volvulus* (nematode) has a same number and length of introns than *T. solium* (cestode), probably because of its close phylogenetic relationship than the other organisms (Table 2). Interestingly, positions of introns and the gene structure in *TsTBP1* are similar to the observed in mouse and human *TBP1*. It shows the importance of this transcription factor for all the organisms, where always the COOH-terminal domain is conserved but not the NH_2_-terminal domain which can bind species specific factors. Figure 3A shows a Southern blot with *T. solium* genomic DNA digested with four restriction enzymes and hybridized with a complete cDNA encoding TsTBP1. It shows a single band with *Hind*III and *EcoR*I, and two bands with *Bgl*II and *BamH*I (enzymes that cut once the TsTBP1 coding region). Figure 3B shows a Northern blot revealing one strong hybridizing transcript of about and 1.1 kbp and a faint transcript about 1.0 kbp, probably degradation of 1.1 kbp band. On the other hand, a quantitative Real-time PCR shows a relative expression of 1.00±0.08 Units for larval and 2.08±0.13Units for adult stages, the mRNA levels of *TsTBP1* were normalized using TsCu/ZnSOD gene (Fig.3C). Southern and Northern blot patterns suggest that *TsTBP1* is a single-copy gene that produces an abundant transcript of ~1.1 kbp, which is double expressed in adult than larval stage. It suggests TBP1 its necessary for growth and production of eggs in the adult stage to continue its life cycle. This finding coincide with higher expression of TBP in the gonads during the gametogenesis observed in rat and mouse during spermatogenesis [25].

**Table 2.**
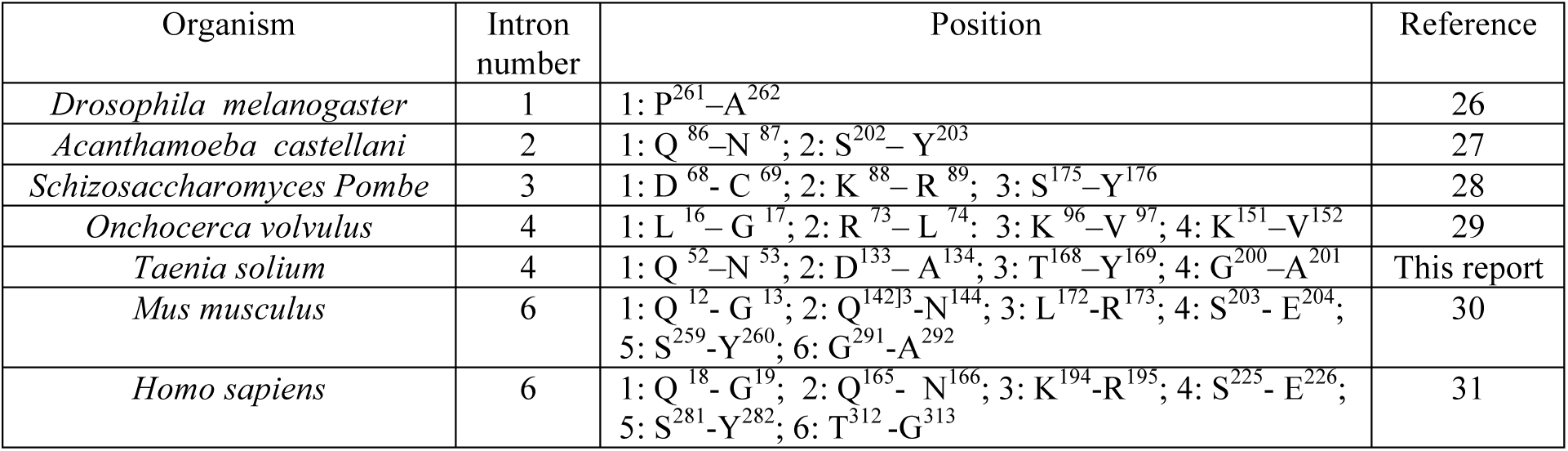
Number of introns in *TBP1* from different species.

**Figure 3.**
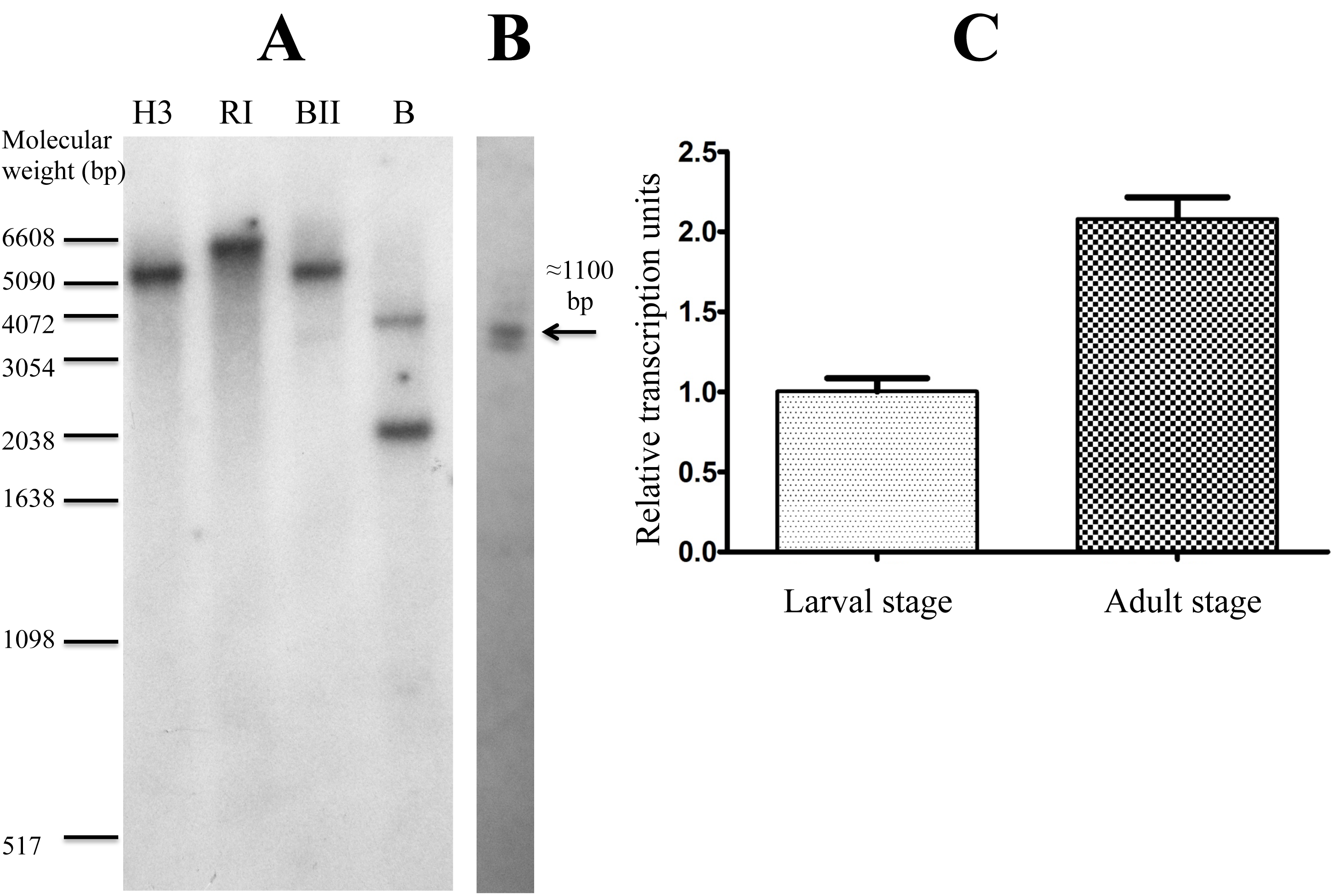
Gene analysis of TsTBP1. A) Southern blot using genomic DNA digested with HindIII (H3), EcoRI (RI), BglII (B2) and BamHI (B). B) Northern blot with total RNA. Both blots were hybridized with a full-length cDNA encoding TsTBP1. Markers of molecular size are placed near of each blot. C) Relative expression of TsTBP1 mRNA in cysticerci and adult stages.

### Identification of TsTAF6 and TsTAF9 in the nucleus of *Taenia crassiceps* cysticerci

Confocal microscopy using DAPI, anti-TAF6 and anti-TAF9 antibodies revealed the presence of DNA, TAF6, and TAF9 in the nucleus of cells that form *T. crassiceps* cysticerci bladder wall. We found that TAF6 and TAF9 are highly localized inside and close to the nuclear membrane. In contrast, a significant but incomplete overlap was observed in the spatial distributions of both factors which suggests a functional transcription (Fig. 4), but places, where these factors are alone it is probably that there are no transcription or incomplete transcription initiation complexes or inhibitory complexes, or are storage sites? Probably, co-localization experiments with RNA Pol II, could help to solve it.

**Figure 4.**
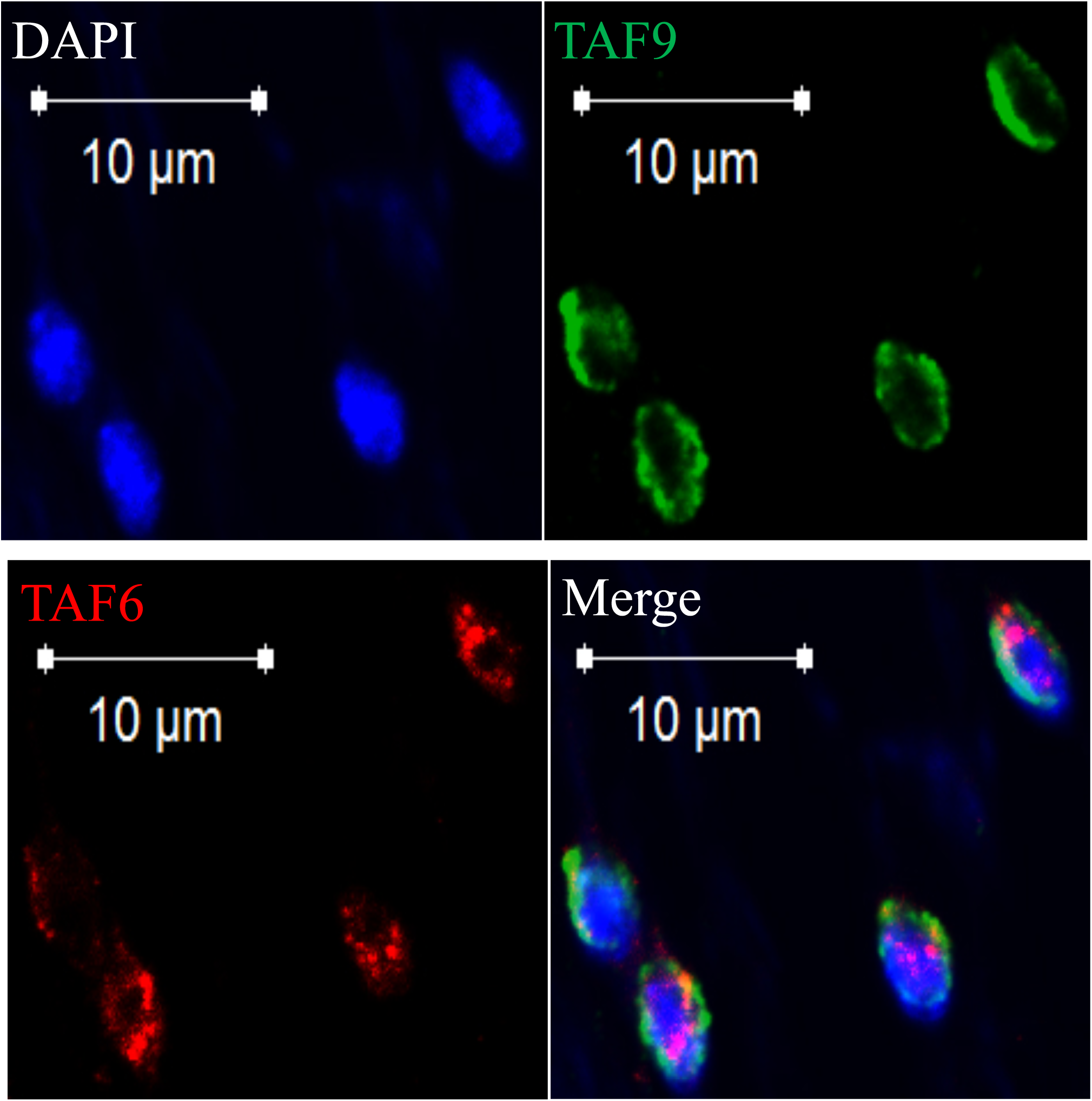
Immunodetection in the nucleus of native TAF6 and TAF9 by confocal microscopy. Taenia crassiceps bladder wall sections were incubated with rabbit anti-TAF6 and mouse anti-TAF9 antibodies and a secondary rabbit or mouse antibody conjugated to Alexa 488 or Alexa 568; in the down-right panel, the merge of the signals produced by both transcription factors is showed.

### Interaction of TsTAF6 and TsTAF9 with DPE of *TsTBP1* promoter

Figure 5 shows the EMSA of *TsTBP1*-DPE. In lane 1 can be observed the free run control; in lane 2, two shifted bands when nuclear extract was incubated with *TsTBP1*-DPE probe, in lane 3 and 4 is denoted the decrease in the intensity of the bands when a homologous competition with 25X and 50X (respectively) *TsTBP1*-DPE cold probe was added to the reaction. In lane 5 and 6, a super-shifted band can be observed when anti-TAF6 or anti-TAF9 antibody was added to the reaction, respectively. In lane 7 we can see the shifted bands produced by nuclear extracts and the consensus DPE probe. Finally, in lane 8 and 9, we observe the disappearance of the bands when consensus DPE and *TsTBP1*-DPE were mutated, respectively. EMSA identify a TGTCGA motif located at +28 to +32, which has a high similarity to consensus DPE (A/G-G-A/T-C/T-G/A/C) and where it is known that TAF6 and TAF9 binds [7, 8, 32]. The DPE is an element mainly present in promoters that lack TATA-box; such is the case of the *TsTBP1*. Interestingly, DPE motif (GCTGT) is found in other *T. solium* core promoter at +30 to +34 bp in pAT5, TsPrx, TsCu/Zn SOD at +29 to +33, and TsTrx1 at +21 to +25. EMSA data shown interaction between TsTAF6 and TsTAF9 with DPE element; suggesting that probably these factors could also bind to the other promoter of the *T. solium* genes before mentioned, however, it’s important to investigate their binding to other genes to establish the functionality of this element. By the other hand, EMSA confirm that putative TATA-boxes in *TsTBP1* located in an unusual position at -97 and -69 bp relative to the TSS, do not bind TsTBP1 (data not shown) contrasting with the binding of it to pAT5 TATA-box gene located -30 relative to TSS [14] which is in agreement with most of the genes with TATA-box, where the position is at -30 to -24 bp [8].

The identification of putative binding sites to TAF6/TAF9 at +27 bp (DPE) and TBP at -97 bp and -69 bp (TATA-box), likewise the detection of TAF6 and TAF9 in nuclear extract of *T. solium* in the *TsTBP1* proximal promoter, but not the TBP, indicate that *TsTBP1* is a TATA-less gene; and it is in agreement with the data that suggest that house-keeping genes typically do not contain TATA-box or Inr, as is the case of the *TsTBP1*. Interestingly, the presence of DPE and TATA-box is not typically observed in the same core promoter, but are present in core promoters of developmental-related genes [33].

**Figure 5.**
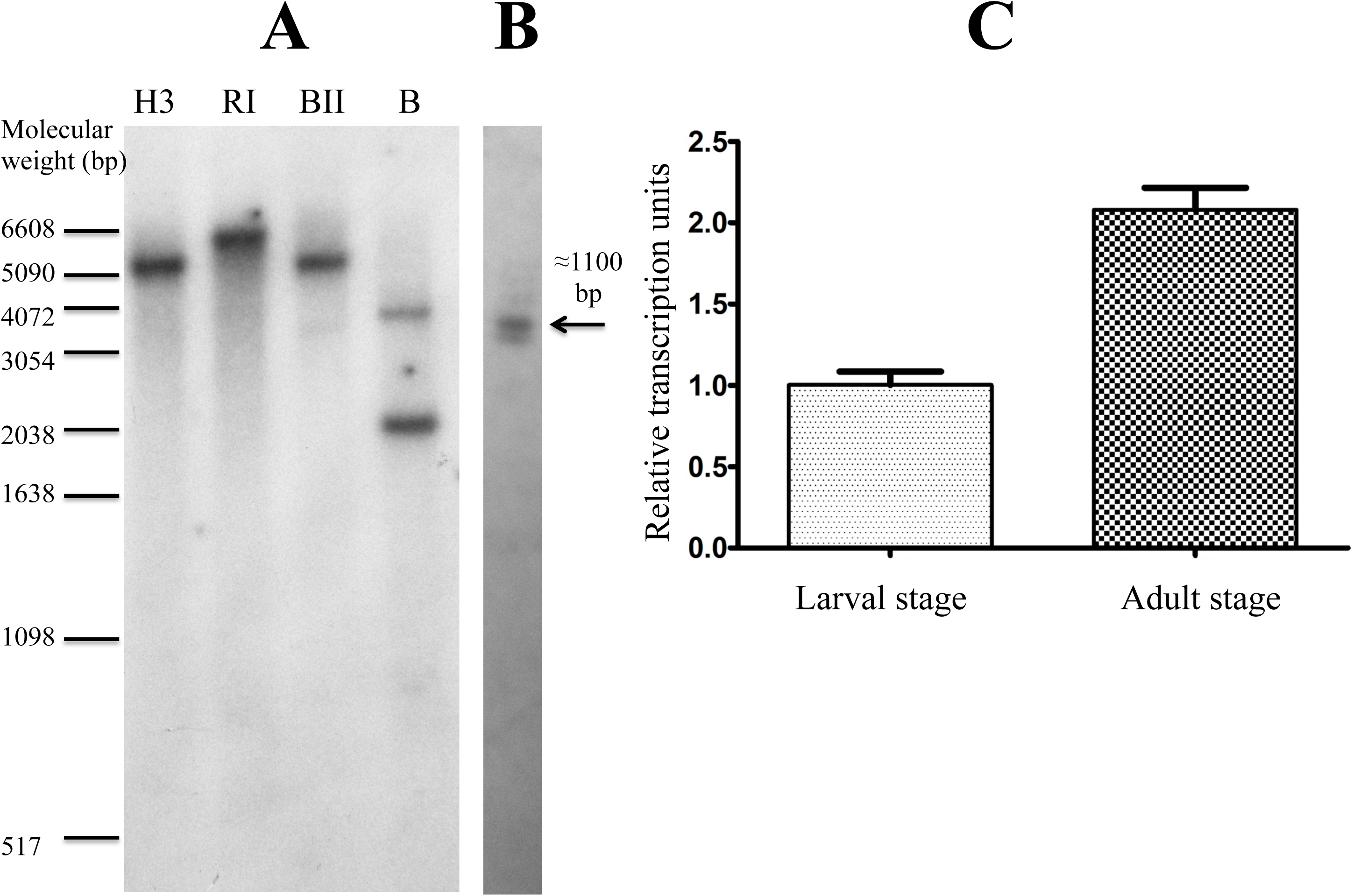
Electrophoretic mobility shift assay showing the interaction of TsTAF6 and TsTAF9 with TsTBP1-DPE. Lane 1: labeled dsDNA-[^32^P] without nuclear extract; lane 2: interaction of native TsTAF6 and TsTAF9 with TsTBP1-DPE probe; lane 3 and 4: competence with TsTBP1-DPE cold probe in a molar excess of 25X and 50X, respectively; lane 5: super-shift interaction using anti-TAF6 antibody; lane 6: super-shift interaction using anti-TAF9 antibody; lane 7: consensus DPE probe interaction with T. solium nuclear extract (positive control); lane 8: mutated consensus DPE probe interaction with T. solium nuclear extract (negative control); lane 9: mutated TsTBP1-DPE probe interaction with T. solium nuclear extract. For probes sequences see Table 1.

### Model of TsTAF6-DPE-TsTAF9 complex

Because, EMSA shows *in vitro* association between two TAFs and the DPE, we cloned and sequenced two cDNA fragments of 1955 pb and 786 pb that encode to *Taenia solium* TAF6 (TsTSF6) and TAF9 (TsTAF9), which show a predicted molecular weight of 70,421 and 29,189 Da, respectively (Fig 6 and 7). To support the experimental association between TAFs and the DPE in EMSA, we studied the complex formation between structural models for TsTAF6 and TsTAF9 and a double helix DNA containing the consensus sequence of DPE (DPE-probe), using molecular modeling.

**Figure 6.**
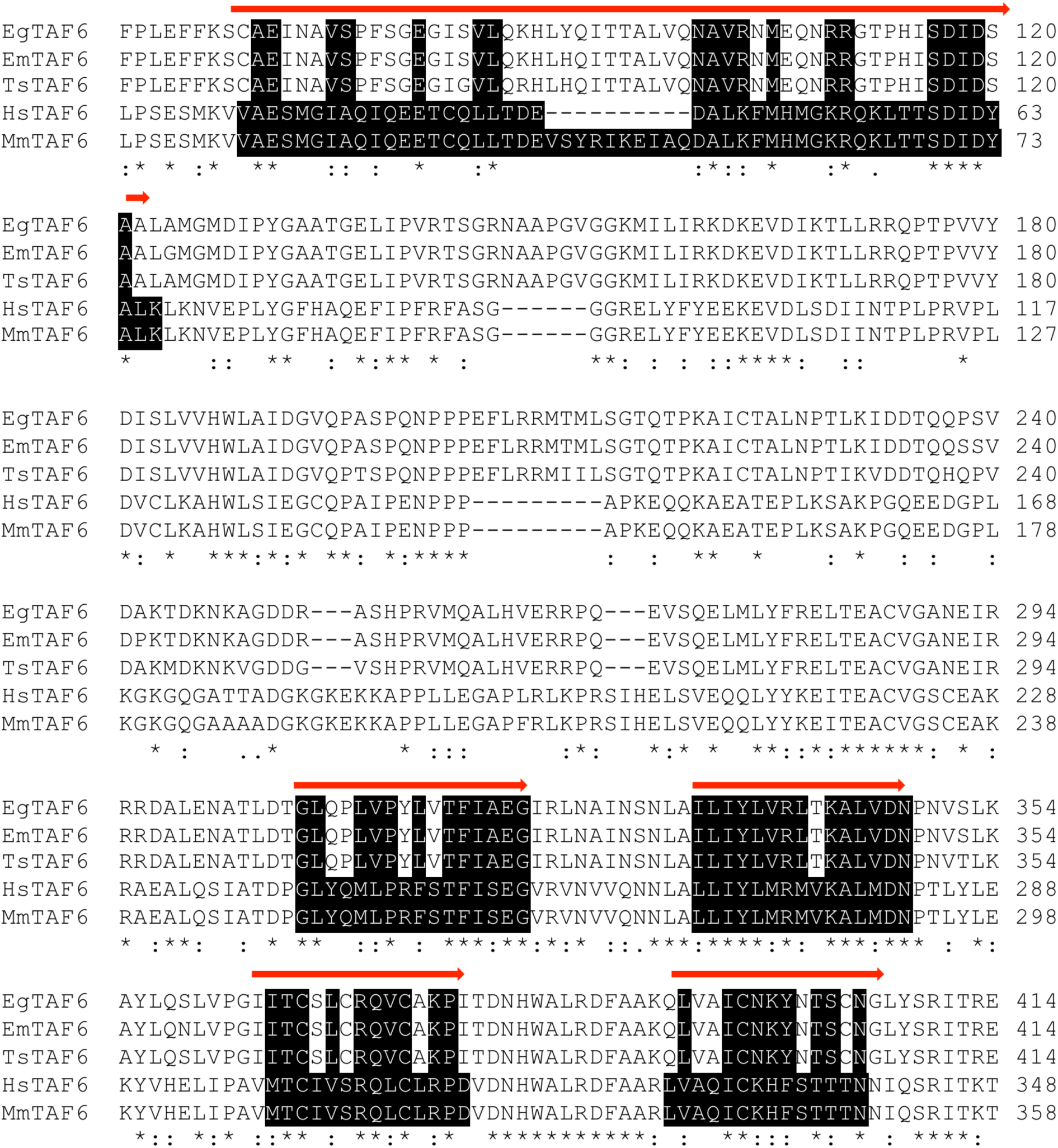
Comparison of deduced amino acids sequences of Taenia solium TAF6 (TsTAF6, KY124274) with others TAF6 from: Echinococcus granulosus (EgTAF6, CDS19795.1), Echinococcus multilocularis (EmTAF6, CDS37851.1), Homo sapiens (HsTAF6, EAL23853.1) and Mus musculus (MmTAF6, AAH58583.1). Arrows represents the Histone Folding domains and the identical aminoacids are highlighted in white letters and black background. Identical amino acids are marked with asterisk (*) and conserved with a colon (:).

It is known human TAF6 structure consists at least of three domains: the histone fold domain (HFD), the middle domain (TAF6M), and a larger C-terminal domain (TAF6C) constituted by several α-helices named HEAT repeats, a motif generally involved in protein-protein interactions. The last two domains form the well-conserved C-terminal region of TAF6 [11]. However, there are no reports of complete 3D complete structure for TAF6 and the secondary structure prediction for very a large sequence of TsTAF6 (645 residues) does not indicate clearly the kind of secondary structure elements presents in the HFD and TAF6M domains, while for the COOH-terminal region propensities for α-helices, β-strands, and random coil are predicted. Particularly, the TsTAF6C domain (from Arg258 to Asp484) is clearly predicted as purely α-helical as like in TAF6C from *A. locustae* (26% sequence identity, PDB ID 4ATG [34]). In our equilibrated final TsTAF6 model (See Material and Methods), the TAF6C domain is formed of five HEAT repeats tightly packed defining a single structural domain, nonetheless the rest of the TsTAF6 (i.e. HFD, TAF6M domains and very C-terminus region) not shown some characteristic fold (Fig 8A). Analysis of the electrostatic potential surface of TsTAF6 shows three positively charged patches: two on the TAF6C domain, and other rather large at the N-terminal region (Fig 8B). This latter characteristic physical signature suggests a specific electrostatic function, maybe a recognition site for DNA. Respect to TsTAF9, prediction of secondary structure consists principally of α-helices with some short strand segments. The final equilibrated modelconsistsofalpha/betafoldandahalfsymmetricalstructurewithnegative electrostatic potential (Fig. 8A).

**Figure 7.**
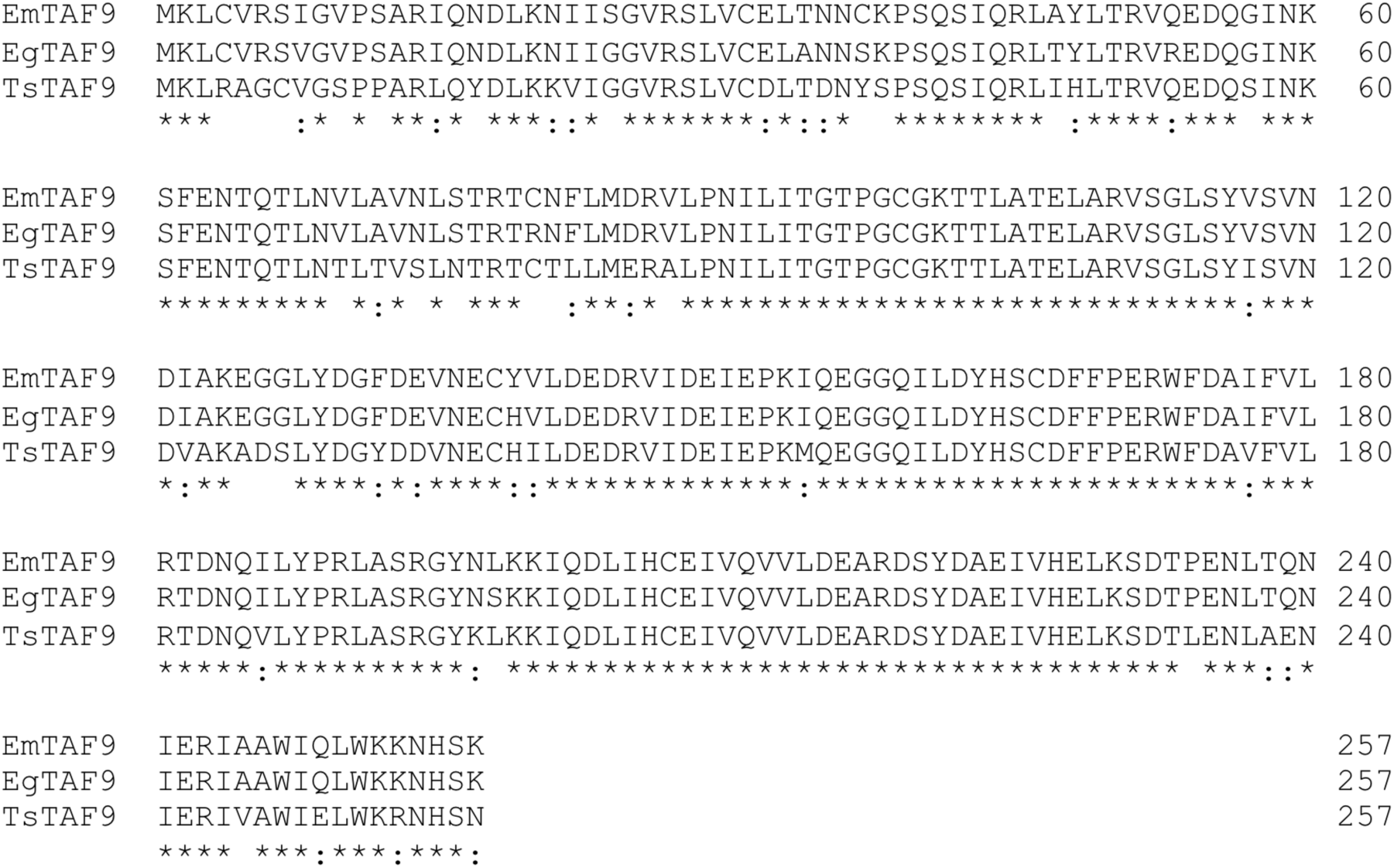
Comparison of deduced amino acids sequences of Taenids’ TAF9. Taenia solium TAF9 (TsTAF9, KY124275); Echinococcus granulosus (EgTAF9, CDS16134.1), Echinococcus multilocularis (EmTAF9, CDS40943.1). Identical amino acids are marked with asterisk (*) and conserved with a colon (:).

**Figure 8.**
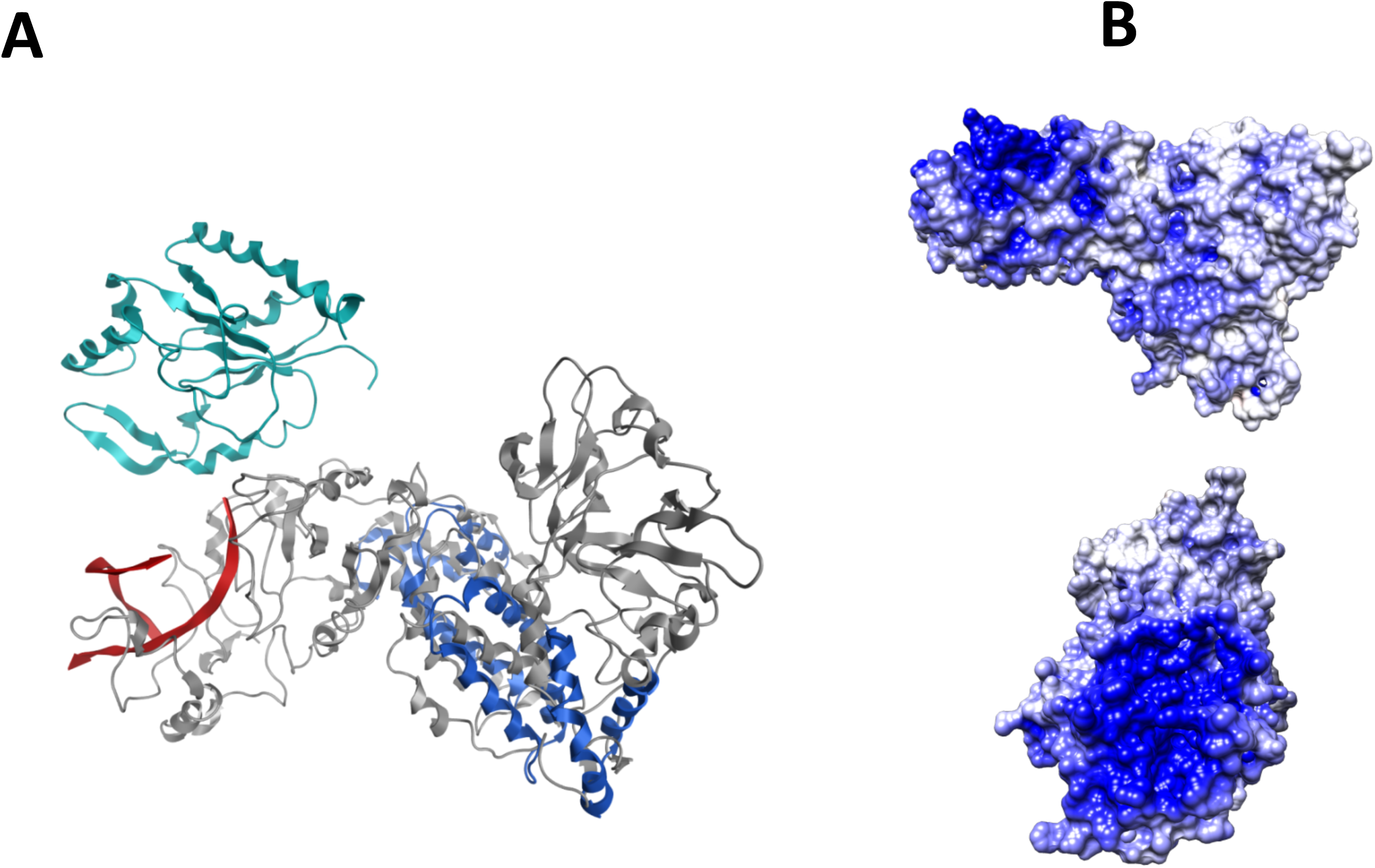
Molecular modeling of TsTAF6 (gray) and TsTAF9 (Cyan) structures. A) The tertiary structure of TsTAF6 superimposed with the five heats repeat from TAF6 of Antonospora locustae (AlTAF6-C, PDB ID: 4ATG, in blue). The RMSD considering only Cα of 195 residues is 7.95 A. B) Representation of the electrostatic potential surface of TsTAF6 (±10 k_B_T/e) showing a prominent positive patch (blue) localized in NH_2_-terminal region of TsTAF6 (lateral and 90° rotation view).

Our study of the association consists of six molecular systems, each containing one molecule of DPE-probe and TsTAF6 inside of a decahedron box (volume: 2528.9 nm^3^). The DPE-probe molecule is placed, in each system, along of the Cartesian axes respect to the center of mass of TAF6C such that any two atoms from different molecules are not closer than 30 Å at the initial configuration. The fact to included only two molecules in the system is supported by the known stoichiometry the high affinity recognition for these systems. When the DPE-probe molecule is positioned facing towards of the positive patch located at the NH_2_-terminal region of TsTAF6 molecule, the formation of the TsTAF6-DPE-probe complex occurred at early time of simulation (5 ns) and the dissociation is not observed for the remaining time of the simulation (120 ns at 300K). The more significant conformational changes upon complex formation included the backbone and side chain of the interacting residues, and eventually changes at the N-terminal region. Such association, do not occur when the starting position of DPE-probe is any of the remainder five system. In this respect, have been suggested that the primary role of the histone fold motif is not be DNA binding but rather a dimerization. and possibly multimerization [34]. In the last frame from MD simulation, complex interactions between double strand of DPE-probe and the TAF6C appear principally mediated by electrostatic and polar interactions involving Arg14, Thr55, Ser56, His58, Arg109, Thr112, Pro113, Thr135 Arg142, Arg146, Arg160 residues and the AGACG nucleotides of DPE (consensus sequence). The re-estimated electrostatic potential of TsTAF9 surface in the complex shows a new landscape, mainly on the TAF6C domain, which now is less positive as a result of the presence of DPE-probe.

We used the last frame from MD simulation of the TsTAF9-DPE system, as a scaffold to assembly the tertiary system TsTAF6–DPE-TsTAF9 by rigid docking. Several of the best scored binding poses in docking studies betweenTsTAF6 and TsTAF9 structures, include those in which TsTAF9 is positioned beside the binding site of DPE-probe, near to conserved HEAT repeats as shown in Figure 7. In fact, mutational experiments suggest that the TAF6C domain modulates the TAF6/TAF9 interactions in *A. locustae* [34]. The contact amino acids residues at the interface region between TAF6/TAF9 in the TsTAF6–DPE-TsTAF9 complex are: Lys8, Lys23, Lys24, Leu26, Leu27, Asn28, Ser29, Arg30, Leu31 and Ala32 respect to TAF6 and Glu60, Ser82, Cys83, Asp84, Asp120, His123, Asp 132, Glu133, Arg 135, Ser137 respect to TAF9. Although TAF9 is not interacting directly with DEP-probe in the consensus model, maybe due to the short in length of the DEP-probe used in the modeling, dissociation of TsTAF6-DPE-TsTAF9 complex is not observed after 70 ns of simulation at 300 K (Fig 9).

**Figure 9.**
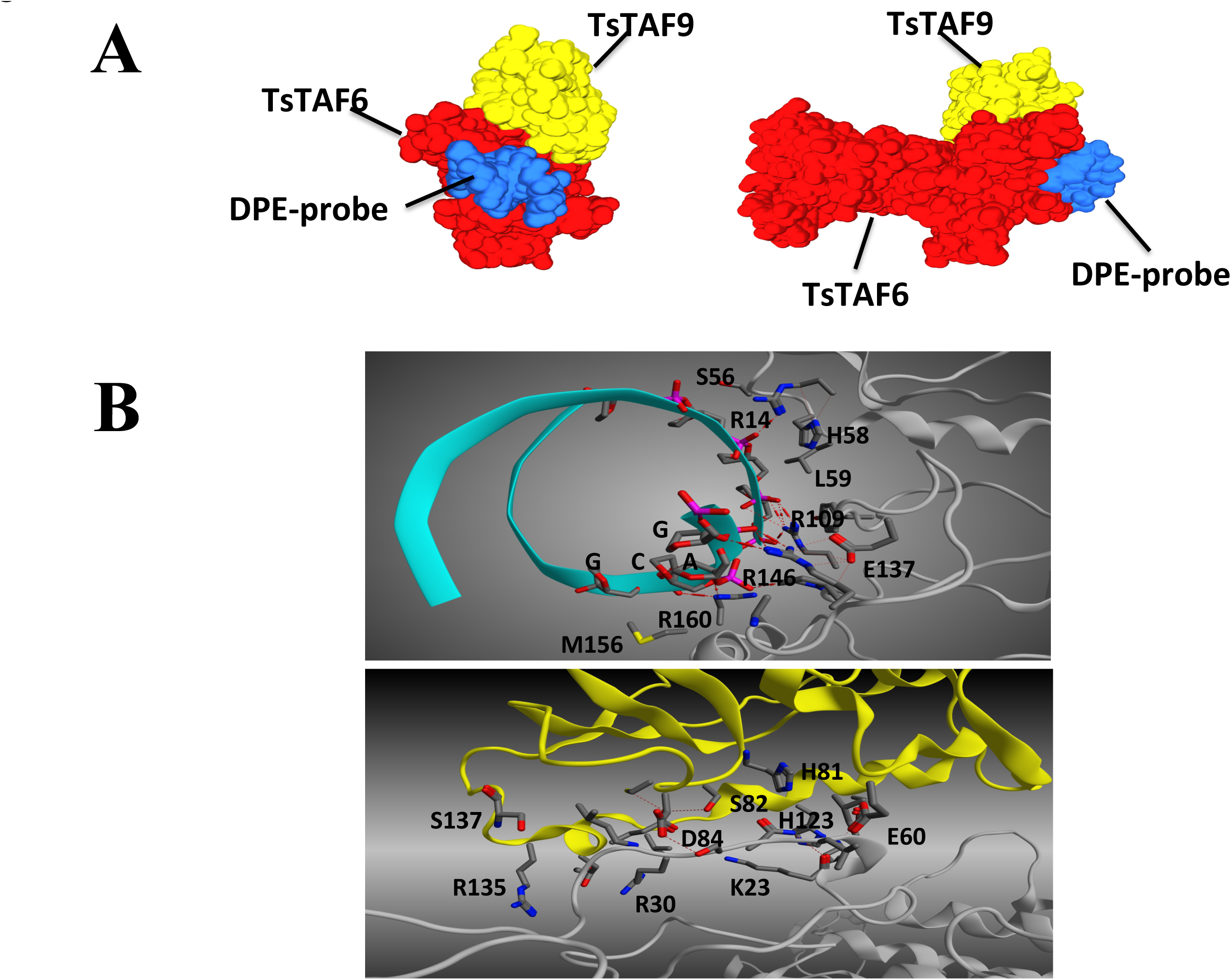
Model of the TsTAF6-DPE-TsTAF9 complex obtained by docking and molecular dynamic simulation. **A)** General view of the final complex in isosurface representation. DPE-probe (blue) is bound to the NH_2_-terminal region of TAF6 (red), while TsTAF9 (yellow) interact with TsTAF6C domain. B) Detail of interaction at 4.5 Å of distance between aminoacid residues and nucleotides TsTAF6-DPE probe (Up) and from TsTAF6-TsTAF9 (Down).

Interestingly, while the alignment of the amino acids deduced complete sequence of TsTAF6 with others TAF6 (Fig. 6) reveals low identity, the HFD domain is conserved between *T. solium*, mouse and human. On the other hand, TsTAF9 also reveal low identity compared to mammals homologous, and as expected, a high identity compared to *E. granulosus* and *E. multilocularis* (Fig. 7). These differences support our hypothesis that TsTAF9 has not a direct contact to DPE, but forms a scaffold for the positioning of the complete TFIID.

### Analysis of core promoter elements presents in *Taeniidae family*

A nucleotide alignment analysis shows two groups the Taeniids promoters genes (Fig. 10A), one TATA-less and other containing the TATA-box in approximately -30 bp relatives to TSS, showing a consensus sequence T-A-T-A-T/A-T/A/G-A/T-G/A/T (TATAWDWD, Fig. 10B) similar to mammalian consensus TATAWAAR [2]. In both groups, the TSS are inside of Inr and corresponds to an adenine (except for TsCu/ZnSOD gene that is cytosine). Inr is widely variable, and a proposed consensus is BBA_+1_NHBN (Fig. 10C), in contrast to the consensus of mammals (YYA_+1_NWYY), the relevant is that most of the TSS correspond to an A_+1_ (8 of 9) (Fig. 4C, [2]). Interestingly, to the DPE shows different forms in core promoter genes of pAT5, TsTrx1 and TsCu/ZnSOD (GGCTGT), TsPrx (TGCTGT), TcPrx (GGTTGC) and in TsTBP1 (TGTCGA), suggesting a consensus G/T-G-C/T-T/C-G-T/A/C similar to the mammalians (Fig. 10D).

**Figure 10.**
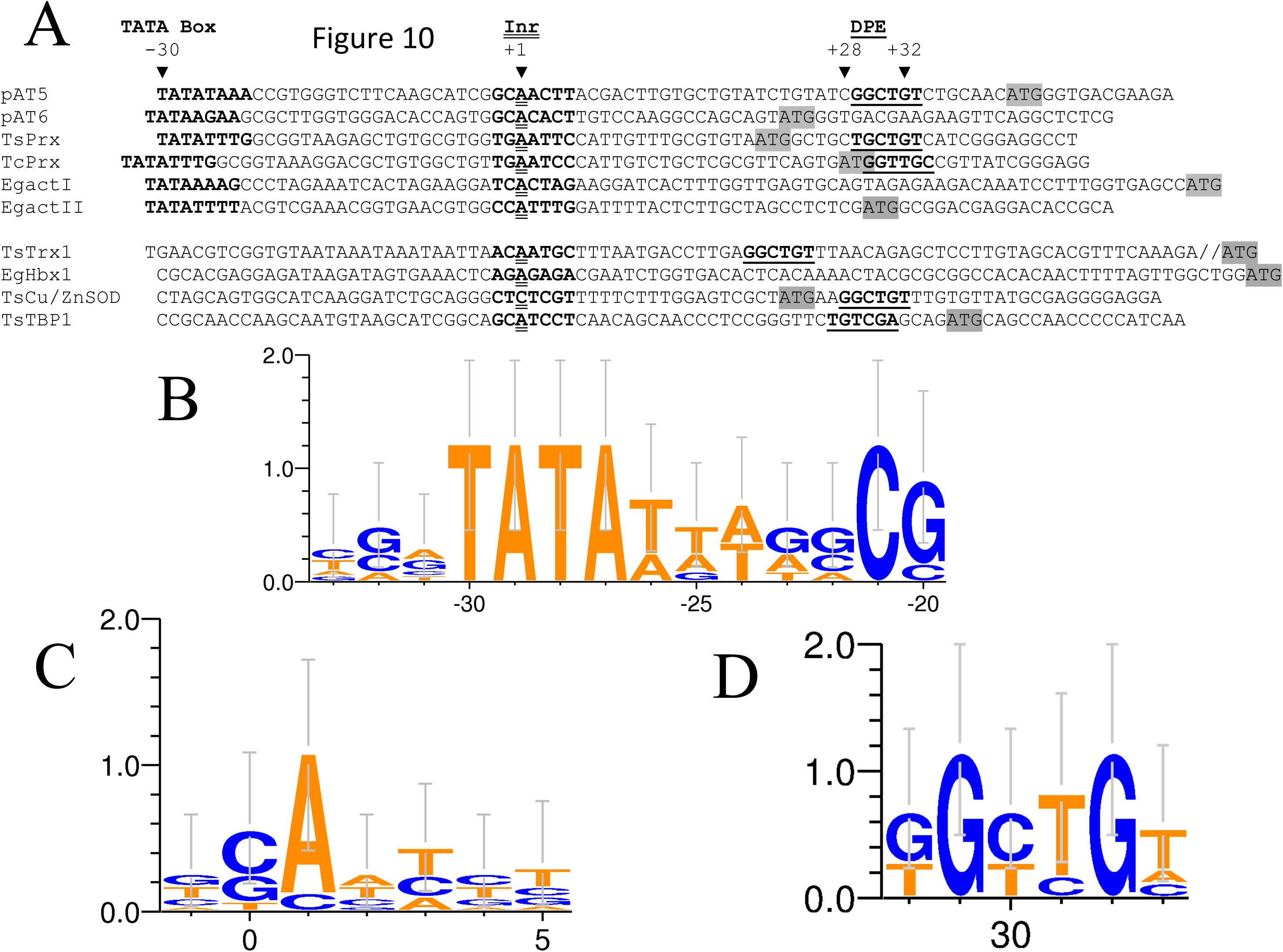
Analysis of regulatory elements presents in core promoters of Taeniids genes. A) Alignment of core promoter sequences from genes of: Actin 5 (pAT5, [35]), Actin 6 (pAT6, [35]), 2-Cys Peroxiredoxin (TsPrx, [35]), Thioredoxin-1 (TsTrx1, [36]), Cu,Zn Superoxide dismutase (TsCu,ZnSOD, [37]) and TBP1 of *T. solium* (TsTBP1); 2-Cys Peroxiredoxin *of Taenia crassiceps* (TcPrx, [35]); and Actin I (EgactI, [38]), Actin II (EgactII, [38]), gen Homebox-containig 1 (EgHbx1, [39]) *of Echinococcus granulosus*. TATA-box in the region approximately of -30 bp (relatives to TSS) is in bold. A_+1_ within the INR is double underlined and the ATG codon is inside a grey box. Putative DPE is in bold and underlined. Consensus sequences to Taeniidae family obtained with WebLogo, for: B) TATA-box, C) INR and D) DPE.

This is the first time that DPE is described to *Taeniids*, and although a low number of core promoters has been characterized to them, the cis-elements found suggest that the regulation of transcription for *Taeniids* includes similar elements than in mammals.

Finally, transcription factors can be activated by physiological, pharmacological and pathological stimuli. Upon activation they exert their physiological effects as described above. In cestodes, transcription regulation remains unknown, but the inhibition of *TsTBP1* or TsTAF6/9 could be a promising target to study gene regulation in *T. solium* and related organisms. This report gives evidence of first TATA-less gene in Taeniidae family and the interaction between the components of the TFIID complex (TBP1, TAF6, TAF9) with cis-elements as TATA-box, and DPE presents in Taennidae family promoters, likewise open new insights in the study of the transcriptional system of this family. In addition, the low identity between TAF9 from cestodes and mammals open the possibility to use it as target to interrupt or modulate the transcription of TATA-less genes in cestodes as a novel therapeutic strategy.

## Acknowledgements

This work was partially supported by Consejo Nacional de Ciencia y Tecnología (CONACyT-176925), and Dirección General de Asuntos del Personal Académico, UNAM (DGAPA-PAPIIT-IN215714). Oscar Rodríguez-Lima was a doctoral student from *Programa de Maestría y Doctorado en Ciencias Bioquímicas* (PMyDCB) *of Universidad Nacional Autónoma de México* (UNAM) and received fellowship from CONACyT-240037.

